# Collecting-Gathering Biophysics of the Blackworm *L. variegatus*

**DOI:** 10.1101/2023.04.28.538726

**Authors:** Harry Tuazon, Chantal Nguyen, Emily Kaufman, Ishant Tiwari, Jessica Bermudez, Darshan Chudasama, Orit Peleg, M. Saad Bhamla

## Abstract

Many organisms exhibit collecting and gathering behaviors as a foraging and survival method. Certain benthic macroinvertebrates are classified as collector-gatherers due to their collection of particulate matter as a food source, such as the aquatic oligochaete *Lumbriculus variegatus* (California blackworms). Blackworms demonstrate the ability to ingest organic and inorganic materials, including microplastics, but previous work has only qualitatively described their possible collecting behaviors for such materials. The mechanism through which blackworms consolidate discrete particles into a larger clumps remains unexplored quantitatively. By analyzing a group of blackworms in a large arena with an aqueous algae solution, we discover that their relative collecting efficiency is proportional to population size. Examining individual blackworms under a microscope reveals that both algae and microplastics physically adhere to the worm’s body due to external mucus secretions, which cause the materials to clump around the worm. We observe that this clumping reduces the worm’s exploration of its environment, potentially due to thigmotaxis. To validate the observed biophysical mechanisms, we create an active polymer model of a worm moving in a field of particulate debris with a short-range attractive force on its body to simulate its adhesive nature. We find that the attractive force increases gathering efficiency. This study offers insights into the mechanisms of collecting-gathering behavior, informing the design of robotic systems, as well as advancing our understanding the ecological impacts of microplastics on benthic invertebrates.

## Introduction

Nature contains many organisms that utilize foraging and gathering behaviors to obtain food. Ants, termites, and bees are all organisms that exhibit collective behavior but have a hierarchical system in place (Frank and Linsenmair 2017; Lemanski et al. 2019; Haifig et al. 2015). Similarly, decorator crabs and assassin bugs are examples of organisms that utilize a gathering behavior to harvest for their survival or hunting, respectively (Thanh et al. 2003; Brandt and Mahsberg 2002). One such organism, the benthic oligochaete *Lumbriculus variegatus*, exhibits this behavior both as an individual or a unified group through the formation of particle clusters. They can gather certain materials in their environments by behaving collectively or individually. Worms are unique in that, unlike ants, termites, or crabs, they exhibit complex, physically-entangled collective behavior without a hierarchy (Ozkan-Aydin et al. 2021; Nguyen et al. 2021a).

Blackworms have been found to actively modify aquatic environments through bioturbation, which involves reworking and ventilation, and their role as biodiffusors and upward conveyors is well-established (Kristensen et al. 2012; Roche et al. 2016). As oligochaetes, blackworms are categorized as a member of the collecting-gathering functional feeding group (FFGs) which harvest and feed on fine particles at and below the sediment-water interface (Cummins and Klug 1979; Ilyashuk 1999; Cook 1969; Wotton 1994). Blackworms use an eversible pharynx to feed and have a mucus-lined body wall that aids in lubrication and respiration, although they may also use their tails for oxygenation (Govedich et al. 2010; Timm and Martin 2015; Tuazon et al. 2022). Though there is no direct evidence for blackworms’ usage of their mucus body wall for collection, it has been shown that particle capture using mucus is utilized by many other macroinvertebrates, such as the polychaete *Chaetopterus*, larval midges (Chironomidae), and the terebellid *Eupolymnia* (Wotton 1994).

Therefore, to investigate the understudied aggregation process of blackworms, we examine their behavior in various settings, including large and small arenas, and through a polymer model of a worm with a short-range attractive force on its body, emulating its sticky mucus layer.

## Materials and methods

### Animals

California blackworms (length 30.2±7.4 mm, diameter 0.6±0.1 mm, mass 7.0±2.4 mg) and algae *(Chlamydomonos reinhardtii*) are obtained from Ward’s Science. Worms are reared in a plastic storage box (35 × 20 × 12 cm) filled with filtered water. They are fed fish food pellets daily, and their water was replaced daily. Worms are kept in water at room-temperature (∼21^°^C) before any experiments. Institutional animal care committee approval was not required for studies with blackworms.

### Data Acquisition

A Logitech Brio 4K webcam (Taiwan, ROC) is utilized to record the large population experiments in a photobox with fixed lighting (∼ 500 lux). Frames are captured in TIFF format using MATLAB’s Image Acquisition toolbox at a rate of 0.20 FPS for two hours, but only the first 90 minutes were used for data analysis.

For the single worm experiments in the small arena, a Leica MZ APO microscope (Heerbrugg, Switzerland) was used with an ImageSource DFK 33UX264 camera (Charlotte, NC) at a frame rate of 30 FPS for one hour with fixed lighting (∼200 lux), but only the first 30 minutes were used for data analysis.

### Data Analysis

Images captured from the webcam are processed using ImageJ software (Schindelin et al. 2012). First, noisy elements are removed from the image by subtracting the background. Then, each stack is binarized with the same color threshold settings to isolate only the green colors corresponding to algae. Finally, the total pixel areas are calculated corresponding to algae over time by analyzing the stack. To obtain the relative amount of algae collected, the data is normalized by dividing each stack by the maximum threshold area of the entire stack and shifting the resulting curve down to zero at the first frame.

Image stacks from the microscope camera are downsampled to 1 FPS using Adobe™ Premiere Pro for analysis. The same protocol as previously described to isolate and analyze the respective colors of materials is followed here. To estimate a worm’s posture, similar thresholding as previously described is performed to segment the worm from the material. A metric called “extent” (*r*_*ee*_) of the worm is defined to estimate the distance between two of the farthest away pixels in the segmented image of a worm. This metric is used to measure how “exploratory” (large *r*_*ee*_) or curled up (lower *r*_*ee*_) the worm is (see Fig. 4 Top). Using this pipeline, the *r*_*ee*_ of a single worm is compared in material versus the same worm in a clean environment.

## Results

### Blackworms’ Collective Gathering of Algae

We aim to explore blackworms’ collecting-gathering behavior, focusing on algae as the material due to its abundance as a food source and distinguishable green color (Fig. 2) (Ilyashuk 1999). We also assess the impact of population size on this behavior. Fig. 1 shows our experimental setup. At the beginning of each trial, we add 10 mg (dry weight) of well-mixed algae to a 10 × 10 cm square petri-dish containing 50 mL of filtered water. We then distribute worms (N = 10, 25, or 50) into the experimental arena. We measure algae mass by drying it using paper towels and feeding the worms beforehand to minimize algae consumption during the experiments. The camera initially detects only a small amount of finely mixed algae. As worms move around the arena, the algae aggregate into a darker green pigment, increasing the relative threshold area. After the aggregation phase, worms consolidate the algae into a singular blob, decreasing the color as they assimilate it and block it from the camera’s view (Fig. 2, bottom). We note that blackworms weakly smell algae and strongly smell each other, as tested using an olfactometer (see SI Fig. 1).

**Fig. 1.**
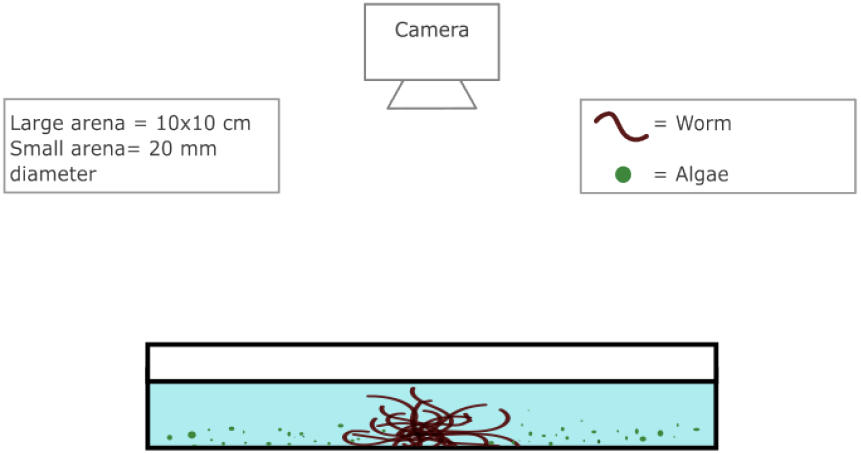
Experimental Setup. Schematic of experimental setup. Large-scale arenas are used for collecting-gathering experiments in a 10×10 cm square petri dish with 50 mL of filtered water and 10 mg (dry weight) of algae. These experiments are filmed from above using a webcam. Small-scale arenas evaluate a single worm’s collecting biophysics using a confocal petri dish with filtered water and 1 mg (dry weight) of material, either algae or microplastics. These experiments are filmed from above using a microscope camera.

**Fig. 2.**
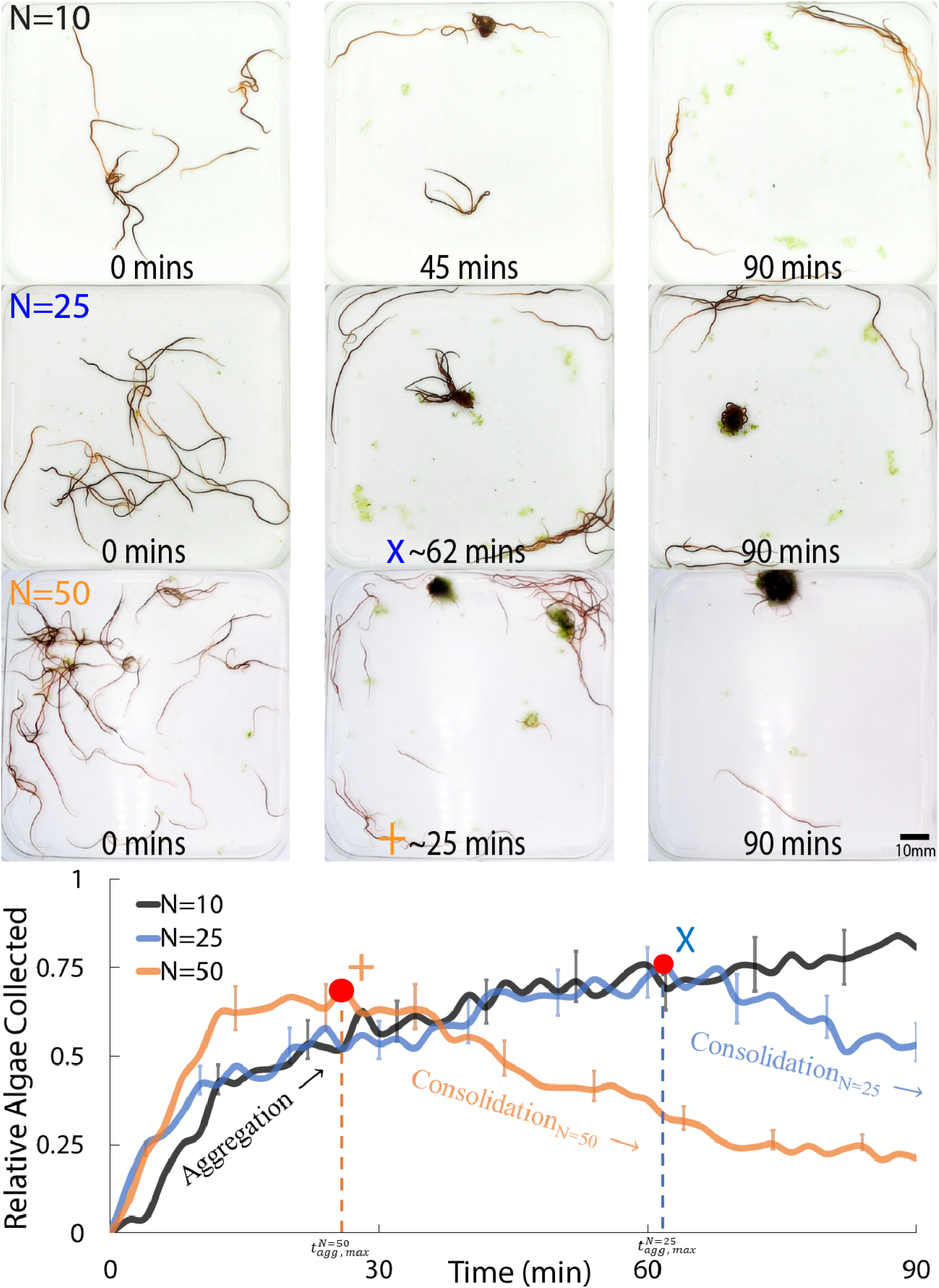
Collective gathering of algae by blackworms. **(Top)** Three blackworm trials with N=10, 25, and 50 worms collecting 10 mg of algae. The maximum aggregation period for N = 25 and 50 are labeled in the graph below. **(Bottom)** Graph showing the relative algae collected for the three populations over time. The data were obtained by thresholding algae in ImageJ. Each curve represents the average relative algae collected, and the vertical bars show the standard error for n=5 trials. (**Supplementary movie 1)**.

Fig. 2 (top) presents instances for three population sizes. The relative algae collection for N = 50 worms (orange curve) reaches a maximum at approximately 25 minutes as they transition from aggregation to consolidation. Decreasing the blackworm population in the arena results in a delayed overall collection of algae, with the time to reach maximum aggregation more than doubling for N = 25 (blue curve) compared to N = 50. This suggests that a larger population of blackworms enhances food resource collection and aggregation, possibly due to social amplification (Amé et al. 2006). Further reducing the population to N = 10 (yellow curve) leads to the failure to reach the consolidation phase within the 90-minute period.

Across these collective gathering experiments, we observe that during the first ∼30 minutes of the aggregation period, the collection is similar for all population levels. On individual worms, the algae appears to adhere to the worm’s body, coalescing into a larger clump as it moves down onto its tail, which may sometimes appear as a ‘hook’ shape (see Fig. 3 algae at t=0,3 mins and supplementary movie 3). Therefore, we next describe the potential mechanisms by which a single worm can aggregate material under a microscope within this period.

**Fig. 3.**
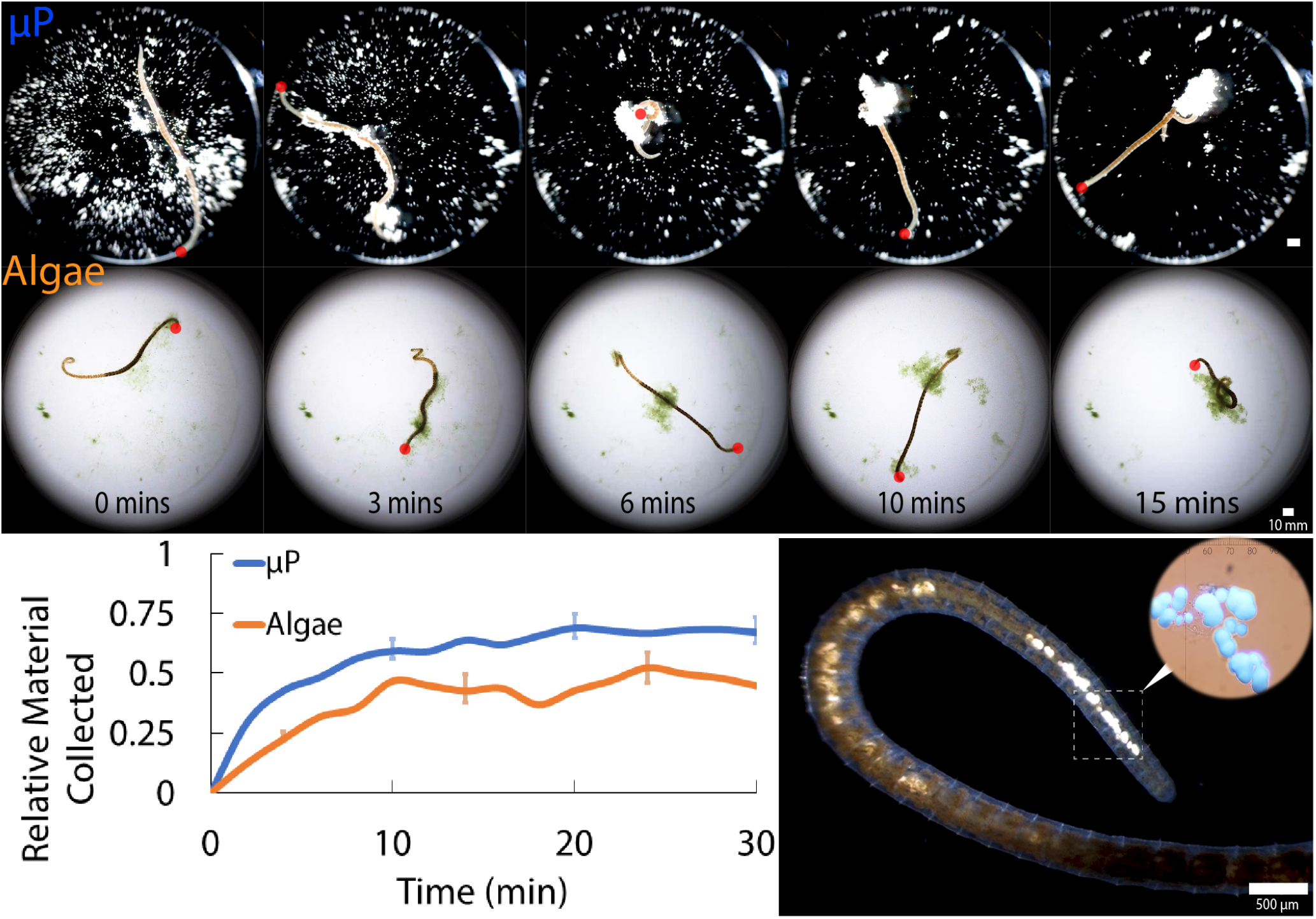
Single worm collecting microplastic materials. Five instances of a worm collecting 1 mg microplastics (*μ*P). The red dot denotes the worm’s head. **(Top)** shows a single worm collecting microplastics (*μ*P, polyethylene microspheres, diameter 68 *±* 6 *μm* and density 1.35g/cc) into one large clump. Microplastics passively adhere to a worm’s body due to externally-secreted mucus. At t=6 mins, the worm passes through a clump on its tail, consolidating particles along its body (**Supplementary movie 2). (Middle)** shows a single worm collecting finely-dispersed algae. A small clump moves downwards along the worm’s body as it moves around (**Supplementary movie 3). (Bottom Left)** Graph showing the relative material collected from *μ*P (blue curve) and algae (orange curve) over 30 minutes. The relative material collected is calculated by thresholding each respective color in ImageJ. Each curve represents the average, with the vertical line showing the standard error for n=5 trials. **Bottom Right** Microplastics inside the digestive tract of a blackworm after several hours of exposure. The ingestion of microplastics can lead to enhanced clumping of particles, as shown in the inset where the excretion of the blackworm results in the formation of a cluster of microplastics. (**Supplementary movie 4)**.

### Single Worm Collecting-Gathering Biophysics

To conduct single worm gathering experiments, we randomly select small, healthy worms (17.6±1.8mm) and place them on the 20 mm glass portion of a 35 mm confocal petri dish containing filtered water and a well-mixed test material. We select smaller worms to prevent interference from the arena walls. Healthy worms are defined as having both anterior and posterior segments that are inspected via a microscope. Algae and microplastics (*μ*P, polyethylene microspheres, diameter 68 ± 6 *μ*m, and density 1.35g/cc) are used as as organic and inorganic materials for the test materials, respectively. Microplastics are chosen as they have more consistent dimensions and are inorganic to test the collection efficiency of indigestible material. Before being exposed to the material, each worm is placed into the arena containing only water for one hour to serve as a control. For each trial, 1 mg of dry material is well-mixed and dispersed into the water before being added to a petri dish.

Fig. 3 illustrates that the collecting efficiency for algae follows a similar pattern to Fig. 2 as the worm packs together the soft material into a 3D spherical shape, which changes the color from light green to dark green. In contrast, microplastics (*μ*P) maintain the same color throughout the trial, resulting in the curve increasing into a steady-state value. The stacks shown in Fig. 3 highlight the dynamics of how a worm aggregates material together. The top and middle panels show that microplastics and algae adhere to the entire length of the worm, coalescing down to its tail. Both materials end up adhering to the worm’s body because of external mucus secretions. One way that a worm may aggregate material along its body is by passing its anterior segments through the clump on its tail, which we refer to as a “threading” aggregation (Fig. 3, top panels from t=3min to 10 mins and supplementary movie 2). Another way is through “peristaltic” aggregation, where, by movement alone, a small clump moves downwards, coalescing any collected particles down to the worm’s tail (Fig. 3, middle panels from t=3min to 10 mins and supplementary movie 3). We observe worms using these two methods regardless of the material.

During the microplastic experiments, blackworms do not consume any of the particles. However, prolonged exposure (hours) results in worms consuming some of the particles which are visible inside their digestive tract (Fig. 3 bottom right). Furthermore, the inset image shows that the blackworm’s excretion results in enhanced microplastic clumping, suggesting that the internal mucus secreted by the worm’s body may also play a role in the aggregation of these particles.

In Fig. 3, we observe that individual worms exhibit relatively less stretched-out movement when the material is in a well-aggregated form around its body (top panel, algae snapshot at 15 mins). To quantify the extent of the worm’s exploration, we measure the largest distance (*r*_*ee*_) between the two furthest pixels on the worm’s body. While the relative material collected shown in Figs. 2 and 3 quantifies the size of the debris cluster, the low-dimensional metric *r*_*ee*_ is a measure to estimate the extent and exploration of the worm itself. The evolution of the metric *r*_*ee*_ to estimate the extent of the worm is plotted in Fig. 4 (top panels) for 30 minutes of dynamics. The black curve in the upper panel is the evolution of *r*_*ee*_ for the case of a worm without any algae present in the petri dish (control), while the blue curve is the same metric in the presence of algae. The videos (see SI movie 2 and 3) show that the worm begins to explore around in the petri dish. This exploration, in addition to sticky mucus around the body, leads to the clumping of these algae around the worm. It is observed that the worm reduces its exploration after this clumping occurs, which is evident in the reduced *r*_*ee*_ in the later stages of the dynamics. Based on previous observations, we hypothesized that the reduction in exploration after clumping occurs is due to thigmotaxis, or the worm’s natural tendency to move towards physical contact with a surface, which in this case is the clumped material on its body. In the lower panel of Fig. 4, the average *r*_*ee*_ of the last 10 minutes of dynamics is plotted for the control and the experiments with debris. This average *r*_*ee*_ value for the control experiment is set to unity for each trial for normalization. We found that the average *r*_*ee*_ reduces substantially (10% - 40% reduction, *n* = 3) when there are algae present in the petri dish. To summarize, our experiments with individual worms show that the uniformly spread particles in the petri dish begin to clump due to the worm’s motion coupled with its sticky mucus on its outer layer. This causes the worm to extend its slender body less and maintain contact with the collected debris. In the next section, we explore the influence of the worm’s adhesive properties on collecting efficiency by utilizing an active polymer model of the worm.

**Fig. 4.**
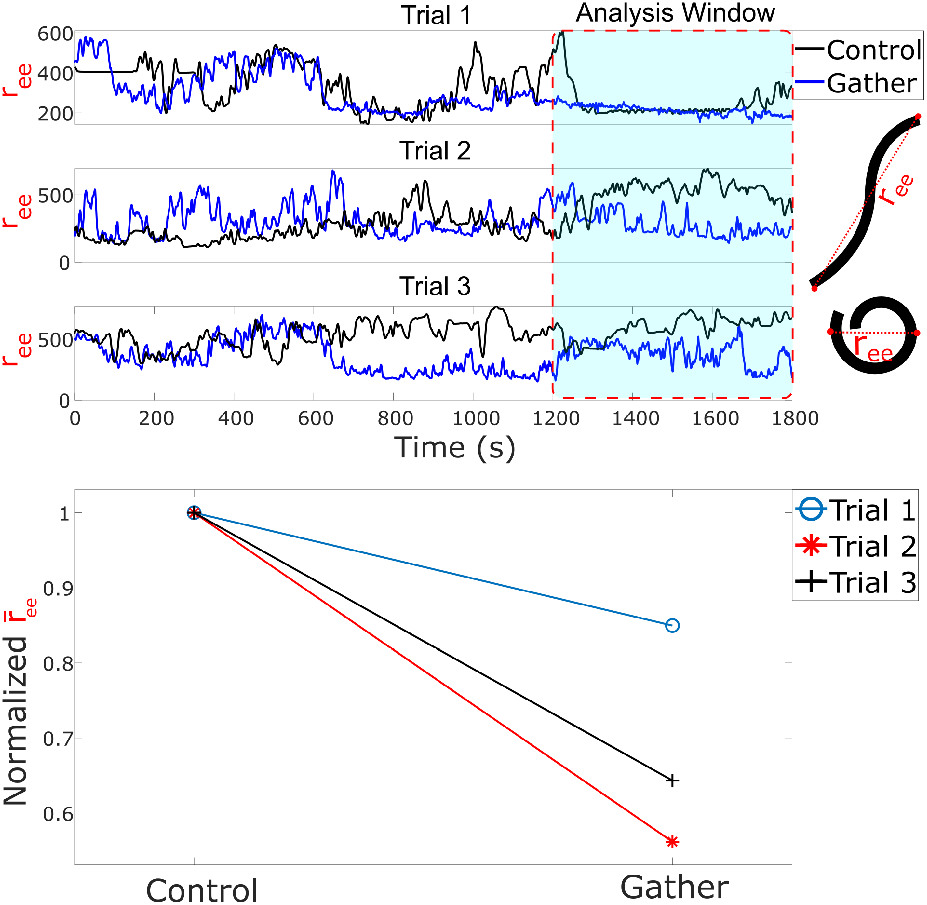
Estimating worm exploration. **(Top)** Timeseries of three different worm trials in 2 scenarios, 1) without any debris (control, in black) and 2) in the presence of well-mixed algae (gather, in blue). **(Bottom)** Average value of *r*_*ee*_ for the last 10-minute window in the top panel, normalized with respect to their respective control values.

### Modeling Collecting-Gathering

Given our observations of worm behavior in collecting and aggregating particles, we develop a simulation model to understand further how worms could accomplish this task. We use an active polymer model of worm dynamics to show that collecting-gathering behavior can emerge from only self-propelled movement and short-range attraction to particles (Fig. 5). Our model is similar to the model described in Nguyen et al. 2021b, in which a worm is represented by a self-propelled active polymer subject to spring, bending, and modified Lennard-Jones-type interaction potentials and uses many of the same parameters, experimentally motivated from single blackworm behavioral assays: number of monomers *N*_*m*_ = 40, equilibrium distance between monomers *σ* = 1.189, single-worm interaction coefficient *ε* = 1, spring constant *k*_*s*_ = 5000, bending stiffness *k*_*b*_ = 10, and self-propulsion force magnitude *F*_active_ = 340. The temperature is set to 0.274 in model units, corresponding to 20^°^C. The worm is constrained to move within a round arena of a diameter 1.14 times the worm’s length, reflecting the parameters of experiments described in this paper. N=100 “hard” (represented by an excluded volume potential) particles of size 0.25*πσ*^2^ simulating microplastics are randomly distributed within this arena. The worm experiences short-range attraction to a given particle, modeled using the same interaction potential governing the worm behavior, but only when the worm is within a short distance (5 particle widths) of a particle and with a higher attraction coefficient *ε*_particle_ = 10. This value is chosen to ensure efficient gathering: too small of an attraction coefficient would result in the worm merely passing by particles without collecting them, and too large of a value would restrict the worm’s locomotion.

**Fig. 5.**
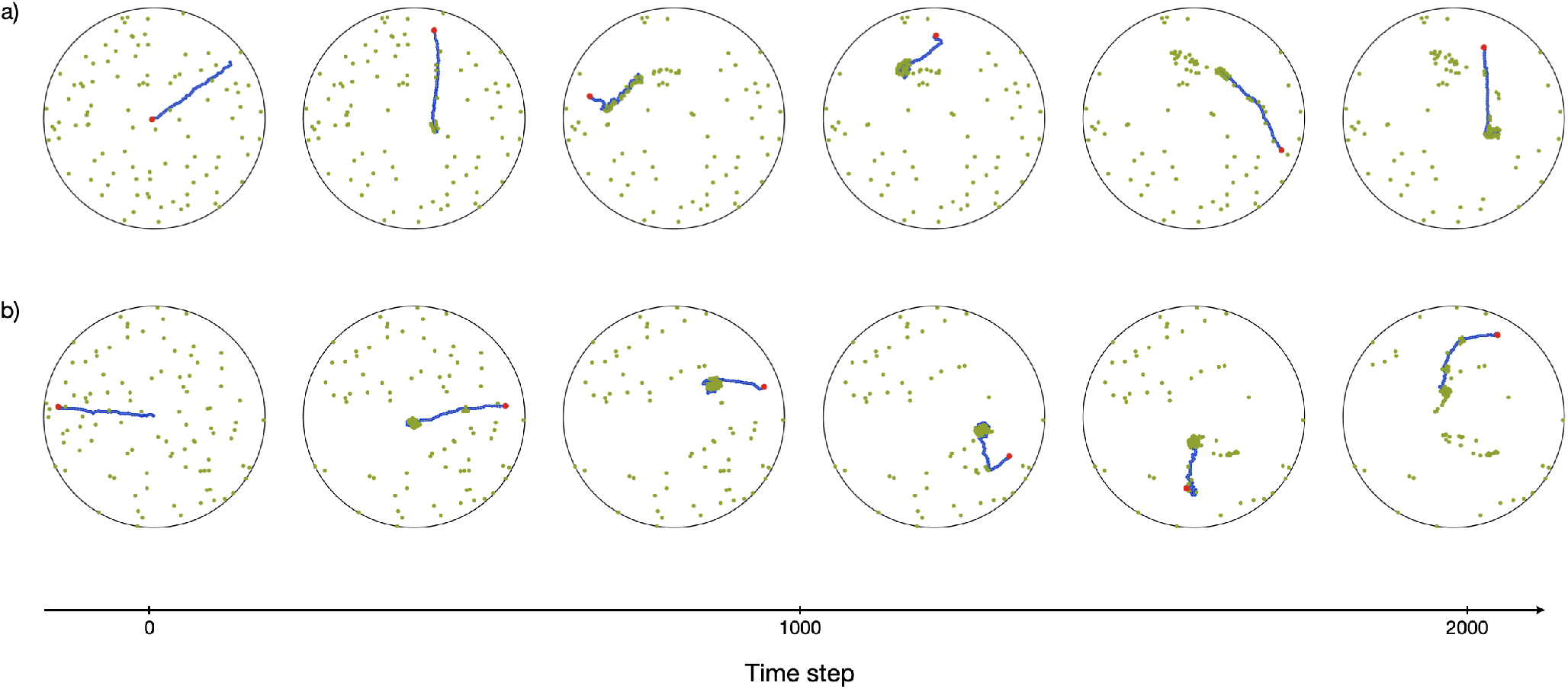
Active polymer worm model of collecting-gathering behavior. We simulate collecting-gathering behavior in an active polymer model of a worm (blue, with a red dot denoting head) governed by self-propelled tangential movement and short-range attraction to particles (green). **(a)** The worm gathers particles into clusters in models without attractive forces between particles and **(b)** with attractive forces between particles, where the attraction represents the worms’ mucosal secretions that bind particles together. **(Supplementary Video 5.)**

We characterize the efficiency of the collecting-gathering behavior by quantifying the average particle cluster area as a function of time. To identify particle clusters, we use the DBSCAN algorithm (Ester et al., 1996) to define a cluster as any group of five or more particles separated by no more than 1 particle width, and we then compute the area of the convex hull of the clusters. Particles not identified as belonging to clusters from this algorithm are assigned to clusters of size 1 particle area. We then determine the average area over all clusters.

We observe that this model is efficient at collecting-gathering particles into clusters. In experiments, we observe that worms secrete a mucosal substance that can bind particles together; to simulate this, we implement attraction between particles, again following a Lennard-Jones-type interaction potential (Fig. 5b and SI movie 5). This attraction results in increased gathering efficiency (Fig. 6), especially at the earlier time steps of the simulation, when the cluster size increases more quickly compared to the model without particle attraction.

**Fig. 6.**
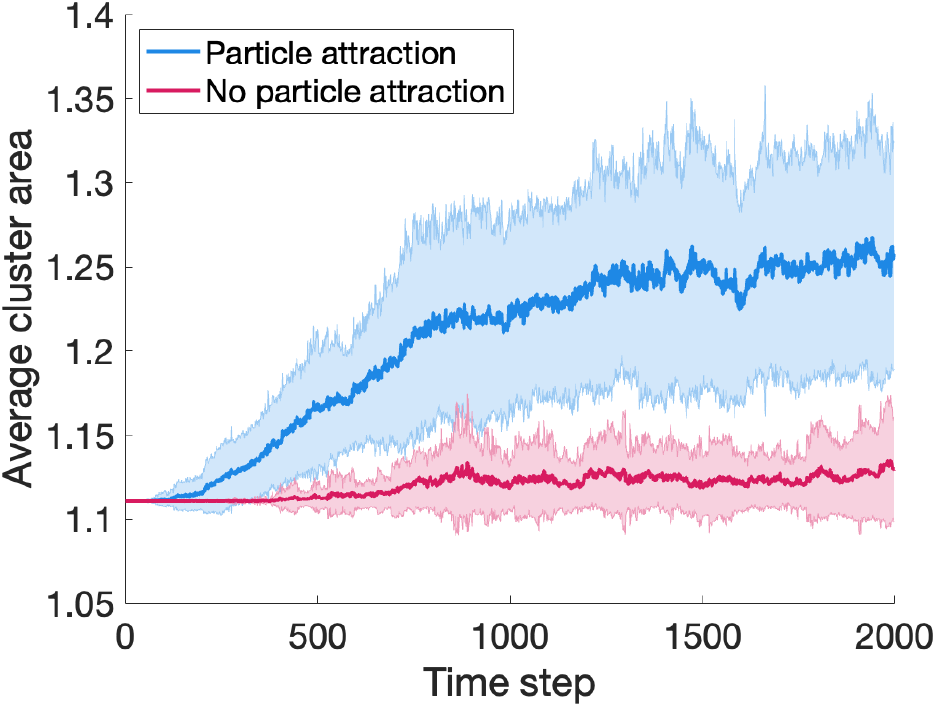
Gathering efficiency in active polymer model. The collecting-gathering efficiency of the active polymer worm is quantified by the average area of particle clusters as a function of time, for the model without particle attraction (pink) and with particle attraction (blue). An average of over 40 simulations is shown for each case, with the shaded regions representing one standard deviation.

## Discussions

Numerous benthic organisms in aquatic environments, including blackworms, modify their surroundings by burrowing and feeding through a process called bioturbation (Kristensen et al. 2012). By burrowing into sediment and keeping their tails above the surface, blackworms ingest sediment from deeper layers and egest it at the surface, making them an upward conveyer. As a biodiffusor, blackworms also alter interfacial sediment through their movement. These behaviors can lead to the formation of significant mounds of sediment (Roche et al. 2016). Blackworms are known to exhibit bioturbating behaviors and are classified as collecting-gathering detritivores based on their feeding behavior. Our results show that the blackworms’ collecting-gathering behavior is influenced by population size, with doubling the population resulting in a reduction of time to reach maximum aggregation by at least half. This increased efficiency could be due to better social and chemical signaling, as blackworms may communicate through olfactory cues (see SI Fig. 1). Finally, as the number of worms in a group increases, there is a greater likelihood of the worms entangling with each other. This leads to increased movement and internal activity within the group, which may facilitate the worms’ ability to collect and gather material from their environment(Patil et al. 2023; Savoie et al. 2023).

By observing the dynamics of individual worms under a microscope, we are able to gain insight into how blackworms aggregate organic and inorganic particles. Our results demonstrate that worms are capable of efficiently collecting particles by using movement and externally-secreted mucus, achieving a single clump for both organic and inorganic materials within roughly 10-15 minutes in a small container.

Furthermore, our observations suggest that the blackworms’ collecting-gathering behavior is not limited to organic matter, as we have also observed them excreting microplastics. Although we did not observe them actively feeding on these particles, this implies that worms consumed microplastics, which is corroborated by other literature that studies their physiological effects on blackworms (Beckingham and Ghosh 2017; Scherer et al. 2017; Klein et al. 2021; Silva et al. 2021).

In nature, as blackworms ingest settled detritus, they could also ingest inorganic material that resides in waterbeds, such as microplastics, which can accumulate significantly at the sediment-water interface of benthic zones in freshwater ecosystems (Krause et al. 2021). Consuming microplastics has previously been shown to cause a severe negative impact on their physiology, such as reduced energy reserves, activation of antioxidants and detoxification mechanisms, and an overall reduction of survival (Klein et al. 2021; Silva et al. 2021). Consequently, the presence and accumulation of microplastics in freshwater ecosystems is of growing concern, as these environments are microplastic retention sites that can transport them downstream to oceans and other bodies of water (Krause et al. 2021). The ingestion of microplastics by benthic macroinvertebrates also raises the concern of microplastic transfer across trophic levels. Benthic macroinvertebrates are eaten by benthivorous fish (Winkelmann et al. 2007), which are consumed by piscivores or larger predators (Vander Zanden and Vadeboncoeur 2002), which could lead to the biomagnification of microplastics along the freshwater food chain. Comparatively, it has been shown in marine environments that it is possible for the trophic transfer of microplastics to occur, as seen in the transfer of microplastics from mussels to crabs (Farrell and Nelson 2013). This is concerning not only to the health of marine and freshwater ecosystems, but to the health of humans that consume animals from these water sources, as microplastics have been recently found in the human bloodstream (Leslie et al. 2022). Though the health risks posed to humans by microplastics have not yet been defined, it is hypothesized that as more microplastics are introduced into the environment and become increasingly bioavailable, health risks to humans will become apparent as they have in a wide range of other species (Koelmans et al. 2022).

### Limitations and Future Outlook

One limitation is that we used was a simplified experimental setup compared to the natural environment of blackworms. For instance, we conducted our experiments in a static system with no water flow, whereas blackworms in the wild are often exposed to flowing water and currents that could affect their feeding and aggregation behavior. Additionally, we used a relatively small number of replicates in our experiments, which could increase the likelihood of random variation influencing our results. However, we found that our experimental system was robust enough to generate consistent results across experiments. Future studies could use larger sample sizes and different experimental conditions to further investigate the behavior of blackworms.

Another limitation of our study is that we focused on only one species of worms, *Lumbriculus variegatus*, which may not be representative of other worm species or other organisms in general. Additionally, our study focused mainly on the physics and biology of the mechanisms underlying blackworm behavior, rather than the ecological or evolutionary implications of this behavior. Future studies could investigate how the gathering-collecting behavior of blackworms affects their survival and reproduction, as well as how this behavior may have evolved over time.

Finally, we acknowledge that our attempt to analyze the worm’s exploration within microplastics was limited by the significant blockage caused by the white particles. Future studies could use alternative methods to evaluate exploration behavior, such as using a different type of particle. Despite these limitations, our study provides valuable insights into the behavior of blackworms and opens up avenues for further research on this unique and complex organism.

## Conclusions

We have investigated the collecting-gathering behavior of blackworms using image analysis and simulations, providing new insights into this functional feeding phenomenon. Our results show that blackworms can efficiently aggregate and consolidate distributed particles into larger clumps using externally-secreted mucus on their bodies and movement. This behavior is influenced by population density and the type of material being collected. Furthermore, our analysis of the extended length of the worm suggests that worms reduce their movement after clumping enough material, potentially due to thigmotaxis. In addition, our simulations have validated the biophysical mechanisms underlying the collecting-gathering behavior of blackworms, demonstrating that this behavior can emerge from self-propelled movement and short-range attraction to particles. Consequently, we also found evidence that blackworms can collect and consume synthetic materials such as microplastics. The idea that blackworms can aggregate materials through ingestion and excretion has been previously explored for sludge reduction, where blackworms have been shown to ingest waste sludge and excrete it as compact feces, decreasing its sludge volume index by half (Elissen et al. 2006). However, the aggregation via excretion of microplastics by blackworms has not been explored to our knowledge.

Overall, our findings have implications for the design of robotic systems inspired by the behavior of blackworms and for understanding the ecological impacts of microplastics. Our study provides new insights into the mechanisms behind collecting-gathering behavior and its potential applications in engineering and environmental science.

## Supplementary data

Supplementary data available at *ICB* online.

## Author contributions statement

H.T. and M.S.B. conceptualized the research. H.T. designed the experiments. H.T., E.K., J.B. and D.C. conducted the experiments, for which H.T., I.T., and J.B. performed the analysis. C.N. performed simulations and modeling. M.S.B. supervised the research. All authors contributed to writing, discussion, and revising the manuscript.

## Acknowledgments

H.T. acknowledges funding support from the NSF graduate research fellowship program (GRFP) and Georgia Tech’s President’s Fellowship. C.N. and O.P. acknowledge funding support from the BioFrontiers Institute at the University of Colorado, Boulder. M.S.B. acknowledges funding support from NIH Grant R35GM142588; NSF Grants MCB-1817334; CMMI-2218382; CAREER IOS-1941933; the Open Philanthropy Project. We thank members of the Bhamla lab for useful discussions and Dr. Emily Weigel for providing expert feedback related to topics of this work. Finally, we thank NSF Aquatic Ecology summer REU for supporting D.C. Text in this paper was revised using ChatGPT-4.

