## Supplementary Document for "Collecting-Gathering Biophysics of the Blackworm *L. variegatus*"

*Integrative and Comparative Biology*, 2023, pp. 1–1

doi: DOI HERE
Advance Access Publication Date: Day Month Year
Symposium

SYMPOSIUM

### Supplementary Information: Collecting-Gathering Biophysics of *L. variegatus*

Harry Tuazon,[1] Chantal Nguyen,[2, ‡] Emily Kaufman,[1, ‡] Ishant Tiwari,[1, ‡] Jessica Bermudez,[1] Darshan Chudasama,[1] Orit Peleg[2,3,4] and M. Saad Bhamla[1,*]

#### Abstract

#### Olfactometer

We conducted studies using a Y-maze olfactometer to uncover chemotactic preferences of the worms when exposed to various substances, including control (no odor), food flakes, plastic (polyester mesh), conspecifics, food pellets, algae, and carbon filter. Fig.1 (top) shows a schematic of the olfactometer which consists of two paths connected to a common leg.

Both upper legs are connected to individual peristaltic pumps which inputs liquid from a beaker containing either water with no odor or water containing another substance. A third pump is installed at the end of the common leg which serves as the drain. All pumps are connected onto an Arduino which control a switch. The switch turns on all pumps for two seconds every minute to provide flow to disrupt diffusion which may confound results.

For each trial, a worm is placed in the middle of the bottom leg and given 15 minutes in the olfactometer. To count as as a decision, a worm's entire body has to pass the imaginary threshold at the base of the leg.

We calculated the proportion of blackworms attracted to each odor source as the number of worms that chose the odor out of the total worms that made a decision, and it is shown as N = worms that decided towards the odor / total worms that actually made a decision (total number of trials) for each material at the bottom of the figure. For the control, the values are expressed as the most-preferred side/any preference (total trials).

Results are analyzed using Chi-square tests per treatment, and the significance of the results are assessed both with and without the Bonferroni correction for multiple comparisons. The proportion data was used to create the graphs in the figure, but the chi-square tests were performed on the count data. Fig. 1 (bottom) shows tabulated results for the olfactometer, all with df = 1. Worms in this olfactometer study displayed clear preferences for food flakes and conspecifics. While the Bonferroni correction revealed that the preference for algae was not as strong as initially suggested, the overall results contribute to our understanding of the factors that influence the worms' behaviors and choices in their environment. The uncorrected p-values indicate significant preferences for food flakes ($p < .001$) and conspecifics ($p < .001$). Algae also showed a significant preference ($p = .0183 < .05$) without the Bonferroni correction. However, after applying the Bonferroni correction ($\alpha = .05/7 = .00714$), only food flakes and conspecifics remained significant. This suggests that worms are highly attracted to food flakes and conspecifics, while their preference for algae may not be as strong as initially indicated.

The standardized residuals from the Chi-square analysis provide further insights into the worms' preferences. Positive standardized residuals greater than 2 indicate a stronger attraction than expected, while negative residuals smaller than -2 imply repulsion. In this study, no results reach the threshold of 2; however, we note that the worms show an attraction to food flakes (1.78) and conspecifics (1.55), and a repulsion towards plastics (-1.88).

#### References

© The Author 2023. Published by Oxford University Press. All rights reserved. For permissions, please

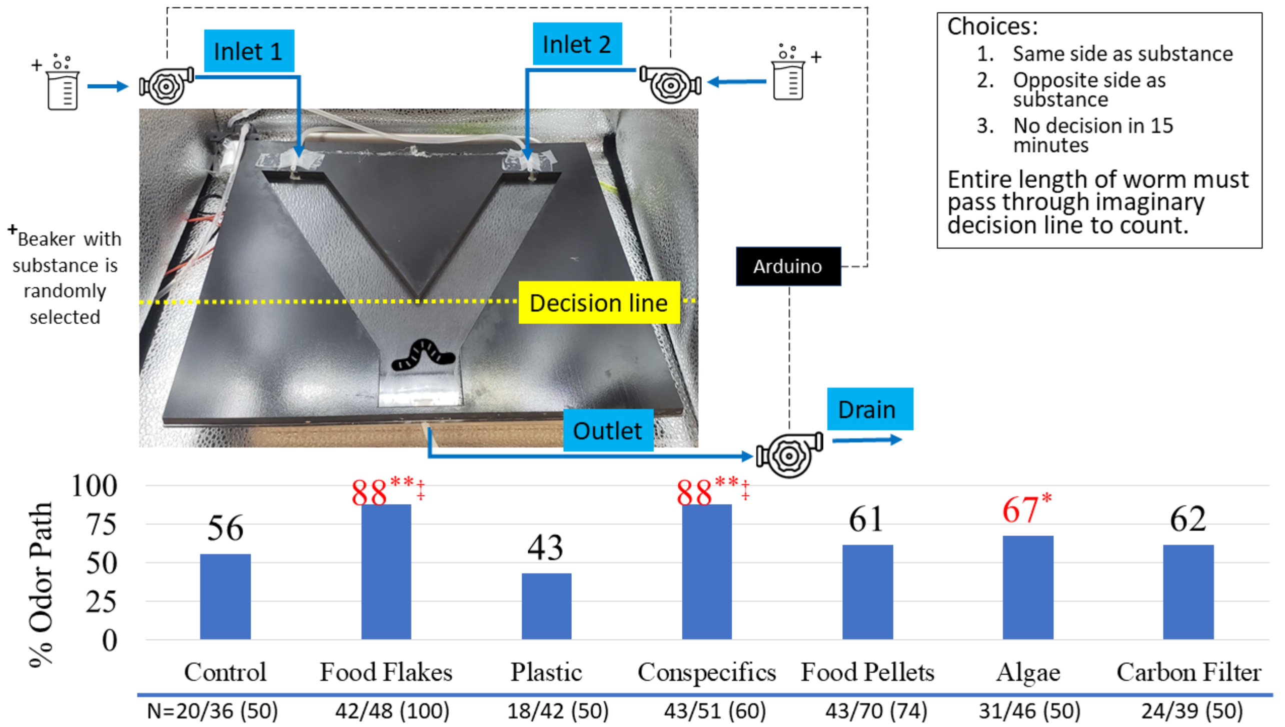


**Fig. 1. Olfactometer test (Top)** displays a schematic of the Y-maze olfactometer used to investigate the chemotactic preferences of blackworms in response to various substances. The olfactometer comprises two paths that are connected to a common leg, with both upper legs linked to individual peristaltic pumps that supply liquid from a beaker containing either water without any odor or water containing a substance of interest. A third pump is positioned at the end of the common leg, which functions as the drain. All pumps are connected to an Arduino, which controls a switch. The switch activates all pumps for two seconds every minute to provide flow and prevent any diffusion that could potentially affect the results. After being placed on the common leg, worms were given 15 minutes to make a decision, during which time the entire length of the worm had to pass through the imaginary decision line in order for it to be counted. **(Bottom)** The chemotactic preference of blackworms was evaluated for control (no odor), food flakes, plastic (polyester mesh), conspecifics, food pellets, algae, and carbon filter. The bar graph shows the proportion of blackworms attracted to the odor source, calculated as N = worms that decided towards the odor / total worms that made a decision (total number of trials) for each odor source, and presented at the bottom of each material.
